# Post-Translational Tubulin Modifications in Differentiated Human Neural Stem Cells

**DOI:** 10.1101/2021.12.31.474563

**Authors:** V. Bleu Knight, Manasi P. Jogalekar, Elba E. Serrano

## Abstract

The tubulin protein fulfills a variety of cellular functions that range from chromosomal separation to locomotion. Functional diversity is achieved through the expression of specific tubulin isotypes in different cell types or developmental time periods. Post-translational modifications (PTMs) of tubulin also are vital for specific intracellular tasks, such as binding and recruiting motor proteins. In neurons, the isotypic expression profile for tubulin is well characterized, and the importance of PTMs for proper neuronal function has gained recent attention due to their implication in neurodegenerative disorders. In contrast, the role of tubulin specializations in the specification of neural cell fate has received minimal attention and studies of tubulin PTMs and isotypes in neuroglia such as astrocytes are relatively few. To bridge this knowledge gap, we undertook an analysis of PTMs in neurons and astrocytes derived from the federally approved H9 hESC-derived human neural stem cell (hNSC) line. In hNSCs, basal cells can be directed to assume neural fate as neurons or astrocytes by specifying different media growth conditions. Immunocytochemical methods, fluorescent antibody probes, and confocal microscopy facilitated image acquisition of fluorescent signals from class III β-tubulin (βIII-tubulin), acetylated tubulin, and polyglutamylated tubulin. Fluorescent probe intensities were assessed with the ‘EBImage’ package for the statistical programming language R, and compared using Student’s t-tests. Qualitative analysis indicated that βIII-tubulin, acetylated tubulin, and polyglutamylated tubulin were expressed to some degree in basal hNSCs and their media-differentiated hNSC neuronal and astroglial progeny. In media-differentiated hNSC astrocyte progeny, quantification and statistical analysis of fluorescence probe intensity showed that acetylated tubulin/ βIII-tubulin ratios were greater than the ratio for polyglutamylated tubulin/ βIII-tubulin. These findings represent a snapshot of the dynamic and varied changes in the tubulin expression profile during the specification of neural cell fate. Results imply that investigations of tubulin PTMs have the potential to advance our understanding of the generation and regeneration of nervous tissue.

## 1. INTRODUCTION

The protein tubulin is abundant in most cells and fulfils specific functions within each cell. Tubulin specializations have been ascribed to isotypic expression profiles and modifications of tubulin that follow protein translation, which are collectively referred to as the ‘tubulin code’ (Janke & Magiera, 2020; Roll-Mecak, 2020). The extensive variety of potential tubulin proteins that can be encoded give rise to the complexity of tubulin functions, which include chromosomal separation, intracellular transport, cellular adhesion and migration, among others (Chakraborti et al., 2016; Seetharaman et al., 2022). PTMs of tubulin, such as acetylation, glutamylation, tyrosination, and glycylation occur enzymatically and serve to confer specific molecular functions (Li & Yang, 2015; Song & Brady, 2015; Pellegrini et al., 2017; Ruse, Chin & Pradhan, 2022; Bär et al., 2022).

Moreover, the alpha and beta isotypic profiles vary according to cell and tissue type, as well as developmental age (Lee, Rebhun & Frankfurter, 1990; Tischfield et al., 2011). The significance of deciphering the tubulin code has been recognized, and many investigations have been undertaken to advance this effort (Cambray-Deakin & Burgoyne, 1987; Lee, Rebhun & Frankfurter, 1990; Audebert et al., 1993; Saragoni, Hernández & Maccioni, 2000; Janke & Kneussel, 2010; Romaniello et al., 2015; Zhang et al., 2015). Translation of the tubulin code is particularly important for central nervous system research because of the putative relationships between posttranslational tubulin modifications and neurodegenerative disorders, including Rett Syndrome, Epilepsy, Alzheimer’s disease, and Parkinson’s disease (Saragoni, Hernández & Maccioni, 2000; Zhang et al., 2015; Delépine et al., 2016; Vu et al., 2017; Bodakuntla et al., 2021; Wu et al., 2022).

Posttranslational tubulin modifications confer specific microtubule specializations in the nervous system. Acetylated neuronal tubulin has been implicated in protein-protein interactions and the trafficking and recruitment of motor proteins depends on the acetylation of microtubules (Pellegrini et al., 2017). Polyglutamylation also affects motor proteins through a demonstrated role in motor protein binding (Lessard et al., 2019). Modification of microtubule associated protein (MAP) is linked to polyglutamylation of tubulin in neurons, and acetylation has a putative role in neuronal plasticity (Moutin et al., 2021). Moreover, the asymmetrical functionalization of microtubules in soma, dendrites, axons, and growth cones through posttranslational glutamylation, acetylation, and tyrosination of microtubules helps establish the architectural polarity that is characteristic of neurons (Park & Roll-Mecak, 2018). Although investigations of posttranslational tubulin modifications in glia are limited, acetylation, and, to a lesser degree, polyglutamylation have been demonstrated in cultured human astrocytes (Knight & Serrano, 2017a). In contrast, while acetylation and polyglutamylation both occur in cultured rat Schwann cells, polyglutamylation is more prevalent in these glial cells (Gadau, 2015). Astrocytic PTMs of tubulin play a putative role in regulation of the Na^+^/ K^+^ ATPase, neurodegeneration, and the inflammatory response (Casale, Previtali & Barra, 2003; Yoshiyama et al., 2003; Youn et al., 2015; Santiago-Mujika et al., 2021). The distinct posttranslational modification profile of tubulin within the nervous system is presumed to contribute to protein function, although a detailed characterization of these relationships is still lacking. In contrast, the isotypic expression profile of tubulin has been studied in greater detail.

The Class III β-tubulin isotype (βIII-tubulin), which is synonymous with neuron specific β-tubulin, has been widely used as a biomarker to identify neuronal differentiation. However, the use of βIII-tubulin in this regard has been scrutinized because its expression is observed in other neural cell types. In particular, βIII-tubulin has been demonstrated in distinct populations of human astrocytes, including astrocytes of fetal origin, as well as mature astrocytes after injury or neoplastic transformation (Katsetos et al., 2001, 2002; Dráberová et al., 2008; Knight & Serrano, 2017b). In H9-derived human neural stem cells (hNSC), the expression of βIII-tubulin increases with the expansion of basal cells, and is observed in cultures grown in neuronal and astroglial differentiation conditions (Oikari et al., 2016). Thus, the isotypic expression pattern of βIII-tubulin appears to intersect the boundary between neurons and astrocytes. The promiscuous expression of βIII-tubulin and other differentiation markers among neural cell types has prompted the exploration of alternative molecules to describe cell fate determination (Oikari et al., 2016). PTMs of tubulin, specifically acetylation and polyglutamylation, are reportedly upregulated in neurons, and therefore offer the potential to enhance the use of βIII-tubulin as a neuronal marker.

While posttranslational tubulin modifications have been explored in the pre-implantation embryo and through the development of knockout mice, there is a paucity of information about the posttranslational modification profiles of tubulin in differentiating stem cells (Houliston & Maro, 1989; Zhang et al., 2008). The exploration of PTMs in differentiating stem cells may reveal specializations of tubulin that occur during the determination of cell fate. To fill this knowledge gap, we evaluated two tubulin PTMs, acetylation and glutamylation, in the H9-derived human neural stem cell line (hNSC) under three media growth conditions: basal, neuronal differentiation, and astroglial differentiation. To enhance reproducibility, our experimental design relied on immunocytochemical methods that implemented antibodies in the Research Resource Identifiers (RRID) registry (Bandrowski et al., 2016). Our findings provide insight into the posttranslational modification profile of tubulin during the specification of neural cell fate in an *in vitro* system.

## 2. METHODS

### 2.1 Cell culture

Gibco® H9 hESC-Derived Human Neural Stem Cells (hNSC; ThermoFisher Scientific, N7800100; lot 1402001; RRID CVCL_IU37) were cultured according to the manufacturer’s specifications. Complete hNSC serum free media (100 ml) comprised 97 mL Knockout DMEM/F-12 (Gibco®, 12660-012), 1 mL GlutaMAX™ (Gibco®, 35050-061), 2 μg bFGF (Gibco®, PH60024), 2 μg EGF (Gibco®, PHG0314), and 2 mL StemPro neural supplement (Gibco®, A10508). Media were sterilized by filtration through a 0.2 μm porous membrane and stored in 20 ml aliquots to avoid reheating.

The T-25 flasks used for basal expansion and the 24- or 96-well plates used for imaging basal cells and differentiated astrocytes were coated with Geltrex™ Reduced Growth Factor Basement Membrane Matrix (Gibco®, 12760; (1:200) for one hour at room temperature. Prior to the addition of cells, Geltrex™ -coated flasks and plates were rinsed with Dulbecco’s modified PBS (DPBS) that included calcium and magnesium, as recommended by the manufacturer. The 24- or 96-well plates used for neuronal differentiation were coated with poly-L-ornithine (Sigma-Aldrich, P3655; 20 μg/mL) overnight at room temperature (∼26°C), then rinsed twice in sterile water before coating with laminin (Gibco®, 23017-015; 10 μg/mL) for two hours at 37°C, 5% CO_2_. Prior to the addition of cells, double ornithine/ laminin coated flasks and plates were rinsed with (DPBS) without added calcium or magnesium, as recommended by the manufacturer.

Frozen ampules of cells were thawed and transferred to sterile 15 ml centrifuge tubes along with 8 ml of media, then centrifuged at 210 G for 3 minutes. All but 1 mL of supernatant was removed to rid cells of cryoprotectant, then cells were resuspended in media and plated into T-25 flasks coated with Geltrex™ (one flask per ampule, ∼8 × 10^4^ cells/ cm^2^) The flasks of hNSCs (passage 0) were incubated at 37°C, 5% CO_2_. In basal cultures, media were replenished every 48 hours that followed. Cells were subcultured by partial digestion when cultures reached 80% confluence according to the manufacturer’s specifications (day 2 for new vials). Briefly, media were removed and cells were rinsed in DPBS without calcium and magnesium before adding 2 mL of pre-warmed StemPRO Accutase (Gibco®, A11105-01) to each flask. Cells were observed for detachment under the microscope, then transferred to centrifuge tubes along with 9 ml of media. Cells were centrifuged at 210G for 3 minutes. All but 1 mL of supernatant were removed, then cells were resuspended by adding an additional 2 ml of prewarmed media and triturated. The first and second passages were used to initiate neuronal and astrocytic differentiation, and for imaging of basal cells. Passaged cells were plated in complete hNSC serum free media, and differentiation was initiated after attachment, between 24 and 48 hours after plating.

Neuronal differentiation was initiated by replacing media with neuronal differentiation media, prepared according to manufacturer’s specifications with 97 mL of Neurobasal® Medium (Gibco®, 21103), 2 mL of B-27® Serum-Free Supplement (Gibco®, 17504), and 1 mL of GlutaMAX™.

Media were sterilized by filtration through a 0.2 μm porous membrane and stored in 20 ml aliquots to avoid reheating. Half media changes replenished cultures every three to four days. If confluence exceeded 75%, differentiated neuronal cultures were passaged, and passage of differentiated neurons was not undertaken past day 7 as recommended by the manufacturer. Differentiated neurons were split 1:2 between 0 and 2 times. Neurons were harvested at day 7, day 10, and day 14. For cultures that proceeded past 7 days, Dibutyryl cAMP (Sigma-Aldrich, D0627) was added for a final concentration of 0.5 mM on days 7, 8, and 9.

Astrocyte differentiation was initiated by replacing media with astrocyte differentiation media, prepared according to manufacturer’s specifications with 97 mL of Dulbecco’s Modified Eagle Medium (ATCC, 30-2002), 1 mL of N-2 Supplement (Gibco®, 17502), 1 mL of fetal bovine serum (ATCC, 30-2020), and 1 mL of GlutaMAX™. Media were sterilized by filtration through a 0.2 μm porous membrane and stored in 20 ml aliquots to avoid reheating. Half media changes replenished the cultures every three to four days. If confluence exceeded 75%, differentiated astrocyte cultures were passaged. Differentiated astrocytes were passaged and divided in a 1:3 ratio between 1 and 4 times during the 21-day course of astrocyte differentiation for all cultures, as recommended by the manufacturer.

### 2.2 Live cell imaging with phase contrast microscopy

Before cells were fixed, Metavue image capture software (Molecular Devices) in conjunction with a Coolsnap HQ CCD camera (Photometrics) was used to capture phase contrast images of live samples. A 20X objective with a correction collar, attached to an inverted Nikon TE-2000 microscope, was used to visualize cells.

### 2.3 Antibody selection

βIII-tubulin was selected as the base target for our comparison of acetylation and polyglutamylation in hNSCs because studies report the expression of βIII-tubulin in basal hNSCs, as well as differentiated neurons and astrocytes (Oikari et al., 2016). βIII-tubulin was detected with rabbit anti-βIII-tubulin antibody (Abcam; catalog # ab52623, RRID:AB_869991). Detection of polyglutamylation and acetylation parallel previous studies (Knight & Serrano, 2017a). However, while the anti-polyglutamylated antibody used in this study came from the same mouse hybridoma clone (B3), it was purchased from a different vendor (Sigma-Aldrich) than the version used in our previous analysis (Abcam). Specifically, glutamylation was assessed with the mouse anti-polyglutamylated tubulin antibody (Sigma-Aldrich; catalog # T9822, RRID:AB_477598) which is reported to detect polyglutamylation of both the α- and β-tubulin isoforms. Acetylation was evaluated with the mouse anti-acetylated tubulin antibody (Sigma-Aldrich; catalog # T7451, RRID:AB_609894) that has been reported to detect the α-isoform. Antibodies were validated with Western blot analysis using protein from Neuronal Human Astrocytes (Lonza, CC-2565) as previously reported in Knight and Serrano (2017a).

### 2.5 Immunocytochemistry

Tubulin in hNSC cultures was labelled with immunocytochemical methods that have been published previously (Knight and Serrano, 2017a), with specific antibody type and dilution as described below. PBST containing 1% Bovine Serum Albumin was used to dilute primary antibodies to the following final concentrations: rabbit anti-class III β-tubulin antibody, 1:300; mouse anti-polyglutamylated tubulin, 1:200; mouse anti-acetylated tubulin, 1:200. The rabbit anti-βIII-tubulin was incubated simultaneously with either mouse anti-acetylated tubulin or mouse anti-polyglutamylated tubulin. Alexa Fluor® 488-conjugated goat anti-rabbit IgG (Abcam; catalog # ab150077, RRID: AB_2630356) and Alexa Fluor® 594-conjugated goat anti-mouse IgG (Abcam; catalog # ab150120, RRID: AB_2631447) were simultaneously diluted at 1:500 in PBS with 1% BSA for secondary antibody exposure. Hoechst 33342 (0.1 μg/ ml) was used to counterstain cell nuclei before mounting slides.

Immunocytochemistry controls were incorporated into the protocol. To determine the specificity of the βIII-tubulin antibody, 1μg/ ml of the βIII-tubulin synthetic immunizing peptide was incubated with the diluted βIII-tubulin antibody for one hour at ambient temperature (∼26°C) before incubating overnight and exposing to Alexa Fluor® 488-conjugated goat anti-rabbit IgG (Abcam; catalog # ab150077, RRID: AB_2630356) secondary antibody. The cross-reactivity of Alexa Fluor® 488-conjugated goat anti-rabbit IgG and Alexa Fluor® 594-conjugated goat anti-mouse IgG were assessed by incubating samples with only rabbit anti-class III β-tubulin antibody (1:300), mouse anti-polyglutamylated tubulin (1:200), or mouse anti-acetylated tubulin (1:200). For the secondary antibody incubation, Alexa Fluor® 594-conjugated goat anti-mouse IgG (1:500) was incubated with samples exposed to the rabbit anti-class III β-tubulin antibody, and Alexa Fluor® 488-conjugated goat anti-rabbit IgG (1:500) was incubated with samples exposed to mouse anti-polyglutamylated tubulin (1:200), or mouse anti-acetylated tubulin (1:200).

### 2.6 Confocal imaging

Confocal images were acquired during multiple sessions using one of two LSM 700 confocal microscopes housed within the UTEP Border Biomedical Research Center (BBRC), depending on availability. Image capture settings were optimized based on control samples for each cell type (basal hNSCs, 7-day differentiated neurons, 10-day differentiated neurons, differentiated astrocytes (21 days) at each session (Table 1). Therefore, a variety of image capture settings and objectives were used. However, images of cells stained with acetylated tubulin and polyglutamylated tubulin were always captured in parallel, with identical settings, to facilitate a comparison of the posttranslational tubulin modifications in the same samples.

**Table 1.** Settings for Confocal Image Acquisition. Image capture settings are reported for each imaging session, along with the number of images and the cell type(s) that were imaged in each session. Settings are given for the respective channels (1, 2, 3) and were consistent for each session. The # of stack images is given for each cell type, and separated by comma.

| Session | 1 | 2 | 3 | 4 | 5 | 6 |
| --- | --- | --- | --- | --- | --- | --- |
| **Laser Power (%)** | 5,5,5 | 5,2.2,5 | 11,5,5 | 5,5,5 | 7, 5, 5 | 13, 7, 11 |
| **Master Gain** | 850, 427, 507 | 850, 529, 529 | 739, 700, 705 | 850, 610, 775 | 850, 610, 775 | 972, 847, 864 |
| **Objective** | 20x, 63x | 20x, 63x | 20x | 10x | 10x | 10x |
| **Digital Gain** | 1 | 1 | 1 | 1 | 1 | 1 |
| **Digital Offset** | 0 | 0 | 0 | 0 | 0 | 0 |
| **Pinhole** | 1 AU | 1AU | 1AU | 1AU | 1AU | 1AU |
| **Cell Type (days in differentiation media)** | Basal, Neuron (7) | Neuron (14) | Astrocytes (21) | Neuron (10) | Astrocytes (21), Neuron (10) | Astrocytes, Neuron (10) |
| **# of Images (acetylated)** | 2, 5 | 2 | 3 | 7 | 5, 5 | 3,2 |
| **# of Images (polyglutamylated)** | 2, 3 | 2 | 3 | 7 | 5, 5 | 3,2 |

Images were captured with an EC Plan-Neofluar 10X/0.3 or EC Plan-Neofluar 20X/0.5 objective mounted on one of two inverted LSM 700 microscopes (Zeiss). Hoechst 33342 was excited with laser λ = 405 emission, Alexa Fluor® 488 was excited with laser λ = 488 emission, and Alexa Fluor® 594 was excited with laser λ = 555 emission. The LSM’s main beamsplitter was configured for 405/488/555/639 wavelengths, with variable dichroic beamsplitters at 579 nm, 587 nm, and 585 nm for tracks 1, 2, and 3, respectively. The SP 555 filter was used to collect emission from the first track (Hoechst), the SP 640 filter was used to collect emission from the second track (Alexa 488), and no filter was used in the third track.

### 2.7 Fluorescence intensity analysis

Fluorescence intensity values were estimated in accordance with previously described methods (Knight & Serrano, 2017a). Maximum intensity projections were exported with a .tif extension using the Zeiss 2009 ZenLightEdition software package. Raw image .tif files were imported and translated into a 2 dimensional array of pixel intensity values between 0 (black) and 1 (white) using a 16-bit range (216 values) using the image processing package ‘EBImage’ for the R statistical programming language (Pau et al., 2010).

The degree of post-translational modification of tubulin (acetylation or polyglutamylation) was estimated as a ratio of the sum of fluorescent signal from acetylated or polyglutamylated tubulin to the sum of fluorescent signal from βIII-tubulin in each image. The ratio of modified tubulin/ βIII-tubulin fluorescent signal was calculated for all maximum intensity projection images from sessions 4 and 5 because of the identical image capture settings for channels 2 and 3, which captured fluorescent intensity from βIII-tubulin and modified tubulin, respectively. Potential differences in mean values of fluorescence intensity ratios were calculated using student’s t-test with significance threshold (α) of 0.01. Comparisons of the ratios of acetylated and polyglutamylated tubulin to βIII-tubulin were made with data acquired from the same cell type (neurons or astrocytes). Comparisons of tubulin acetylation between neurons and astrocytes were made using the ratios of acetylated tubulin to βIII-tubulin for both cell types. Comparisons of tubulin polyglutamylation between neurons and astrocytes were made using the ratios of polyglutamylated tubulin to βIII-tubulin for both cell types. Statistical power was calculated as the ability to reject the null hypothesis (1-β) using G*Power version 3.1.9.2 (Faul et al., 2009).

### 2.8 Figure preparation

The Zeiss 2009 ZenLightEdition software package was used to manipulate the contribution of the fluorescence signals from each channel to visualize all signals in the merged confocal images, and to export confocal images as TIFF files. The image adjustment settings were consistent for all images in each figure and did not affect the data that were used for quantification purposes. Alignment, labelling, and assembly of images were completed with Adobe CS6 Photoshop.

### 2.9 Rigor and reproducibility practices

Rigor and reproducibility guidelines proposed by the NIH were used as a model for these experiments (Landis et al., 2012). The hNSC line was derived from the WA09 (H9) embryonic stem cell line (NIH approval number NIHhESC-10-0062). The NIH registry for human embryonic stem cells retains the information about the WA09 (H9) stem cell line from which the hNSC line was derived. The federally approved H9 (WA09)-derived embryonic human neural stem cell line (NIH approval number NIHhESC-10-0062) was de-identified by the NIH and as such is exempt from review by the NMSU Institutional Review Board.

hNSC differentiation was initiated within 3 passages (10 population doublings) to guarantee lineage according to the vendor specifications. The human origin of the cell line was confirmed through the alignment of the hNSC transcriptome to the human genome in RNA-seq experiments described previously (Knight & Serrano, 2017c). βIII-tubulin was detected with rabbit anti-βIII-tubulin antibody (Abcam; catalog # ab52623, RRID:AB_869991) that was validated through western blot analysis and by blocking with the immunizing peptide, as described previously (Knight & Serrano, 2017a).

## 3. RESULTS

### 3.1 Basal stem cells and their media-differentiated neuronal and astroglial hNSC progeny are characterized by morphological differences

Phase contrast images of basal and differentiated hNSCs revealed differences in cell size and structure. The basal stem cells (Fig. 1 A-C) appeared smaller in size than the differentiated astrocytes (Fig 1 D-F). Astrocytes appeared to branch and spread out in comparison to hNSCs, which demonstrated a compact morphology. Astrocytes were maintained in lower density cultures due to the tendency for endogenous growth factors to impede differentiation, and the necessity for those same growth factors to maintain stem cell pluripotency and prevent differentiation. Images of hNSC cultures (Figs. 1 and 2) reflect the recommended seeding density of 50,000 cells/ cm^2^ for basal stem cells, and 25,000 cells/ cm^2^ for astroglial differentiation, as well as the passage of astrocytes throughout differentiation when 80% confluence was attained. Despite the high seeding density of 50,000 cells/ cm^2^, differentiated neurons were sparse in comparison to the basal hNSC cultures.

**Figure 1.**
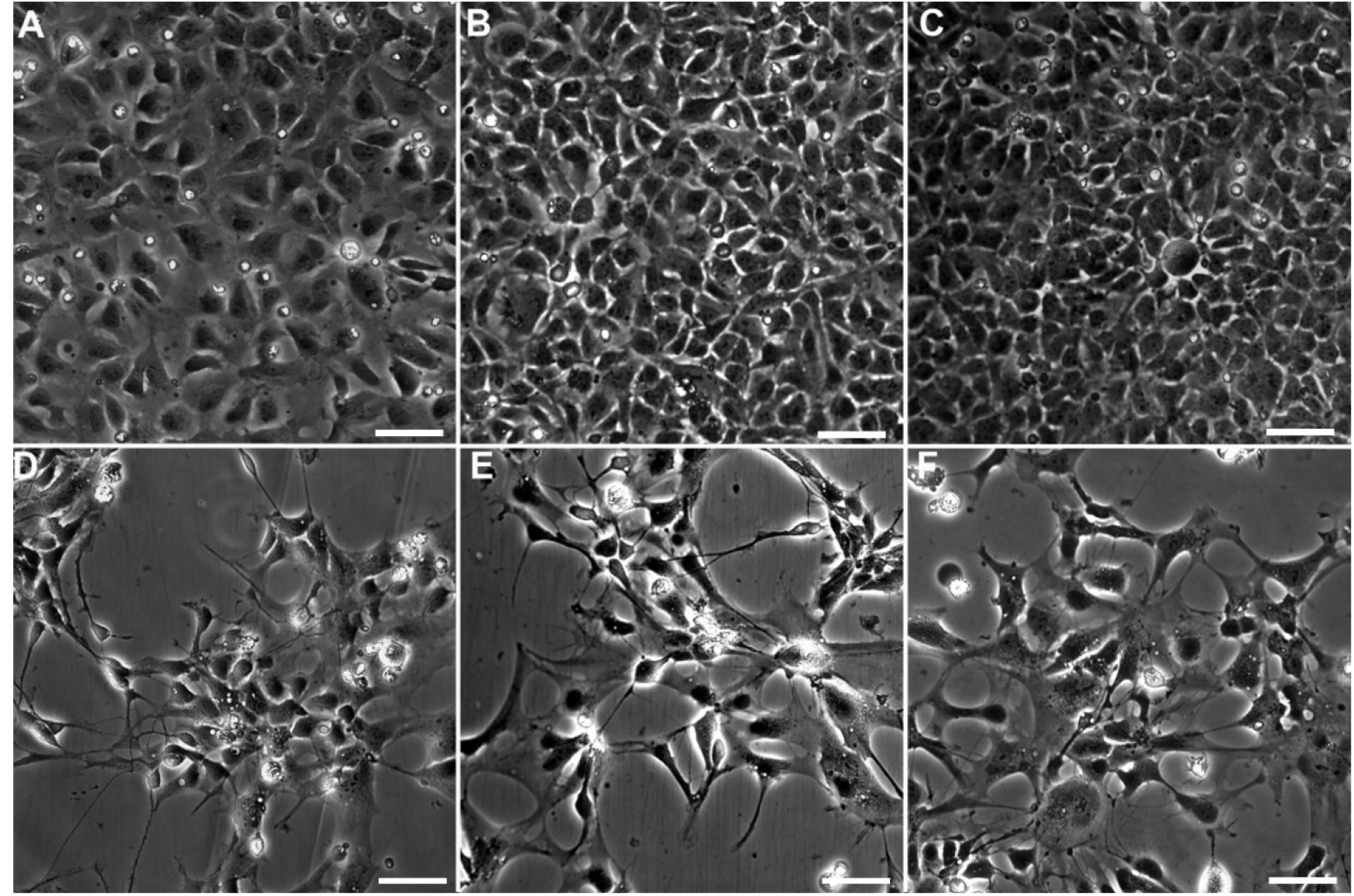
Basal hNSCs and Differentiated Astrocytes. Phase contrast images of live cells depict phenotypic differences between basal hNSCs 3 days after plating (A – C) and astrocytes exposed to differentiation conditions for 21 days (D – F). Scale bar = 50 μm.

**Figure 2.**
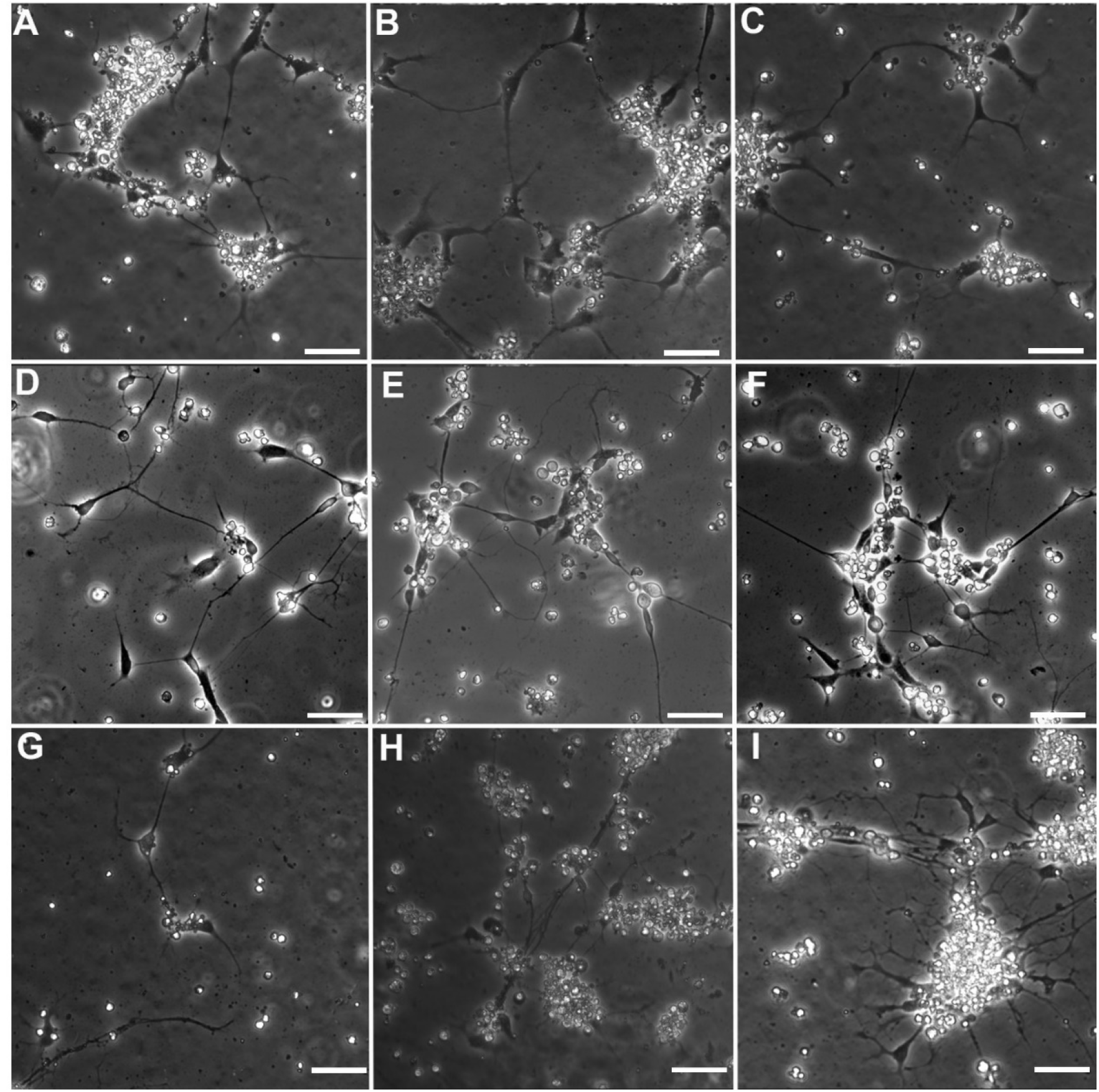
hNSCs in Neuronal Differentiation Conditions. (1 – 4). Phase contrast images of live cells depict increased branching and polarization as neuronal differentiation progressed from day 7 (A -C) to day 10 (D-F) and day 14 (G-I). Phase bright cell bodies and polarization became apparent by day 10 (D-F). While polarization continued through 14 days of differentiation (G-I), phase bright cell bodies were less apparent in cultures after 14 days of differentiation conditions, and debris became prevalent. Scale bar = 50 μm.

Neuronal differentiation was assessed after 7 days, prior to the addition of cAMP (Fig. 2 A - C). The cultures were exposed to cAMP on days 7, 8, and 9, and assessment was undertaken on day 10 (Fig. 2 D-F), and day 14 (Fig. 2 G-I). The rounding up of hNSCs was seen after 7 days and became more prevalent as cultures progressed.

Morphological assessments of basal and differentiated hNSCs were undertaken with phase contrast microscopy (Figs. 1 and 2). Basal hNSCs mandate a high seeding density and grow rapidly, thus they are densely packed and relatively small in comparison with differentiated astrocytes (Fig. 1). After exposure to astrocyte differentiation media for 21 days, many cells become stellate, and some flatten out and spread to resemble the shape of fried eggs (Fig. 1). A qualitative assessment of neuronal differentiation does not show similar changes in the size of the cell body; however, long extensions protrude out of the cell body that become more extensive as differentiation proceeds (Fig. 2). After 7 days, the polarized phenotype that is typical of neurons was not always apparent in differentiated neurons (Fig. 2 A-C) In contrast, after 10 days, some cells appeared to adopt a polarized morphology, with a single dominant axon emerging from one side of the cell body (Fig. 2 D-F). After 14 days, debris became prevalent in the culture and phase-bright cell bodies were less apparent than in the cultures exposed to differentiation conditions for 10 days (Fig. 2 G-I). After 7 days, some cell bodies of differentiated neurons were phase bright, and while branching was evident, a qualitative assessment revealed that branching was not as extensive as compared to differentiated neurons after 10 and 14 days of exposure to neuronal differentiation conditions.

Our assessment of morphology at 7 days, 10 days, and 14 days after the initiation of neuronal differentiation led to a detailed, quantitative analysis of neuronal-differentiated cultures at day 10. We noted that although neurons and basal hNSCs are plated at the same initial seeding density, the cultures maintained in neuronal differentiation media begin to show signs of cell death and debris over time. Ten days following the initiation of neuronal differentiation, hNSC cultures assumed morphologies strongly resembling neurons, and retained the phase bright cell bodies that are indicative of healthy neuronal cultures.

### 3.2 Qualitative immunocytochemical characterization indicates expression of βIII-tubulin and tubulin PTM in basal stem cells and their media-differentiated neuronal and astroglial hNSC progeny

βIII-tubulin expression and the occurrence of post-translational acetylation and glutamylation in hNSC cultures were evaluated with immunocytochemical methods. A positive label for βIII-tubulin was observed in basal hNSCs and in cells cultured under all differentiation conditions (Fig. 3-9). This result is congruent with previous reports of expression of βIII-tubulin in hNSCs, human fetal astrocytes, and cultured neurons (Dráberová et al., 2008; Oikari et al., 2016; Knight & Serrano, 2017a). βIII-tubulin expression was consistently seen in hNSCs after 10 days of neuronal-differentiation as well as after astrocytic differentiation. A qualitative assessment of βIII-tubulin revealed an increase in expression as differentiation in basal hNSC cultures progressed to the astroglial or neuronal state.

**Figure 3.**
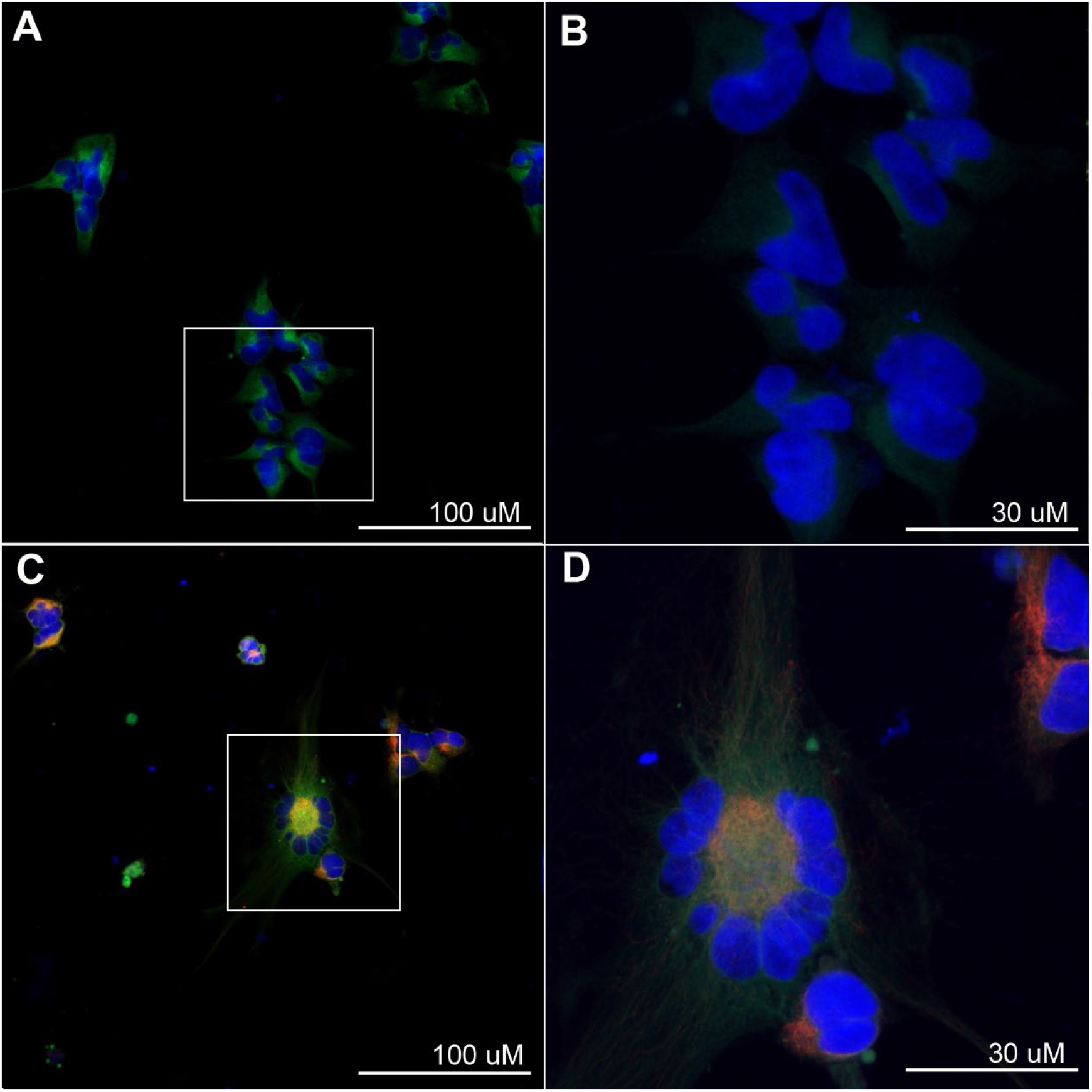
Acetylated, Polyglutamylated and βIII-tubulins in Basal hNSCs. Fluorescent signals from nuclei (blue), βIII-tubulin (green), and acetylated (red, A, B) or polyglutamylated (red, C,D) tubulin were captured from basal (undifferentiated) hNSCs in single optical sections at 20x magnification (A, C) and z-series confocal stacks at 63x magnification (projections; B, D). Regions of image capture from higher magnification images are depicted in white boxes in A and C. Image contrast was enhanced to display fluorescent βIII-tubulin signal (85% contrast), polyglutamylated tubulin signal (A,B; 93% contrast), and acetylated tubulin signal (C,D; 93% contrast). Image capture settings and enhancements paralleled those for images of hNSCs after 7 days of neuronal differentiation (Figs. Fig. 4 and Fig. 5).

**Figure 4.**
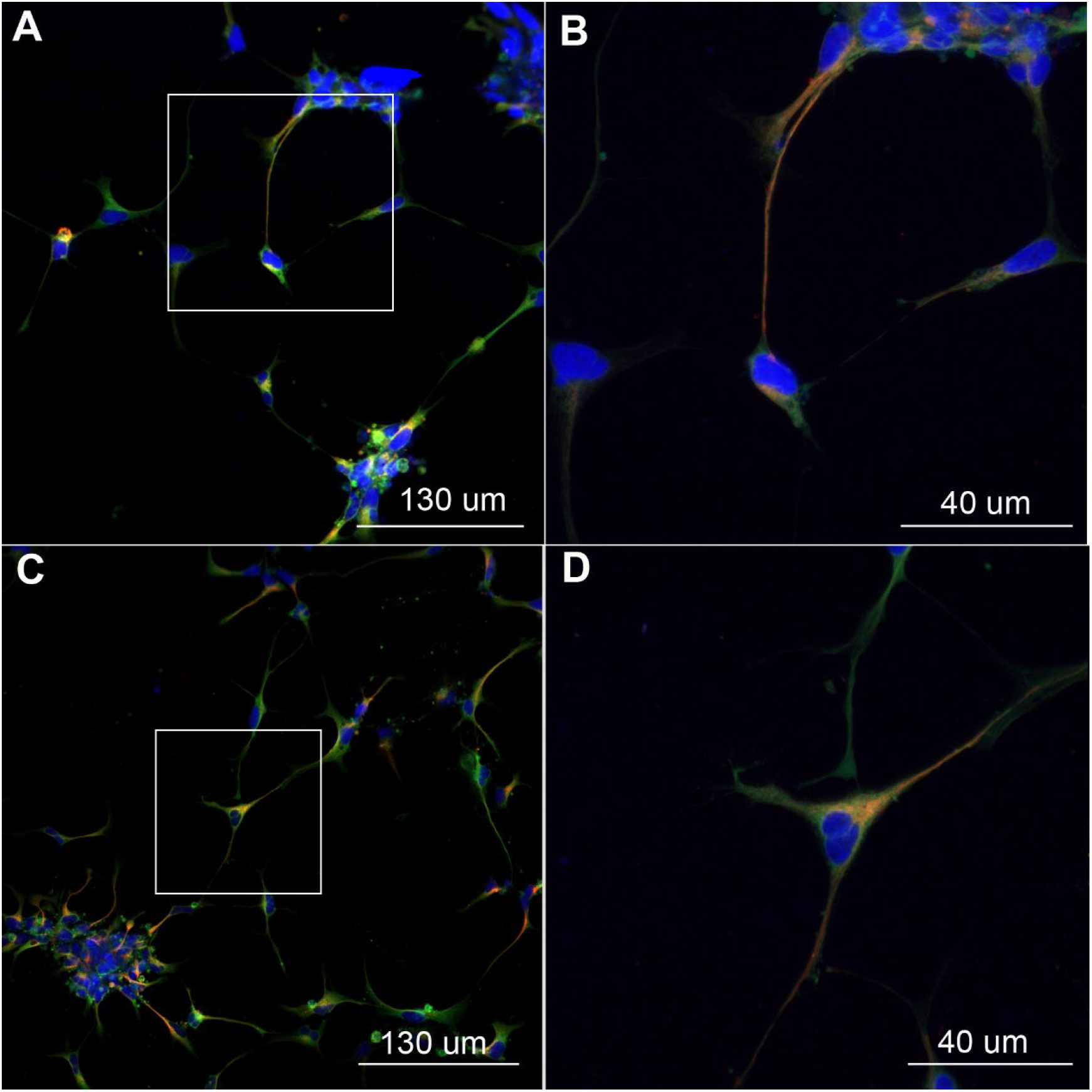
Acetylated Tubulin and βIII-tubulin in Differentiated Neurons (7 day) Fluorescent signals from nuclei (blue), βIII-tubulin (green), and acetylated (red, A, B) or polyglutamylated (red, C,D) tubulin were captured from hNSCs after 7 days of neuronal differentiation in single optical sections at 20x magnification (A, C) and z-series confocal stacks at 63x magnification (projections; B, D). Regions of image capture for higher magnification images are depicted in white boxes in A and C. Image contrast was enhanced to display fluorescent βIII-tubulin signal (75% contrast), and acetylated tubulin signal (93% contrast). Image capture settings and enhancements paralleled those for images of acetylated and polyglutamylated tubulin in basal hNSCs (Fig. 3), and after 7 days of neuronal differentiation (Fig. 5).

**Figure 5.**
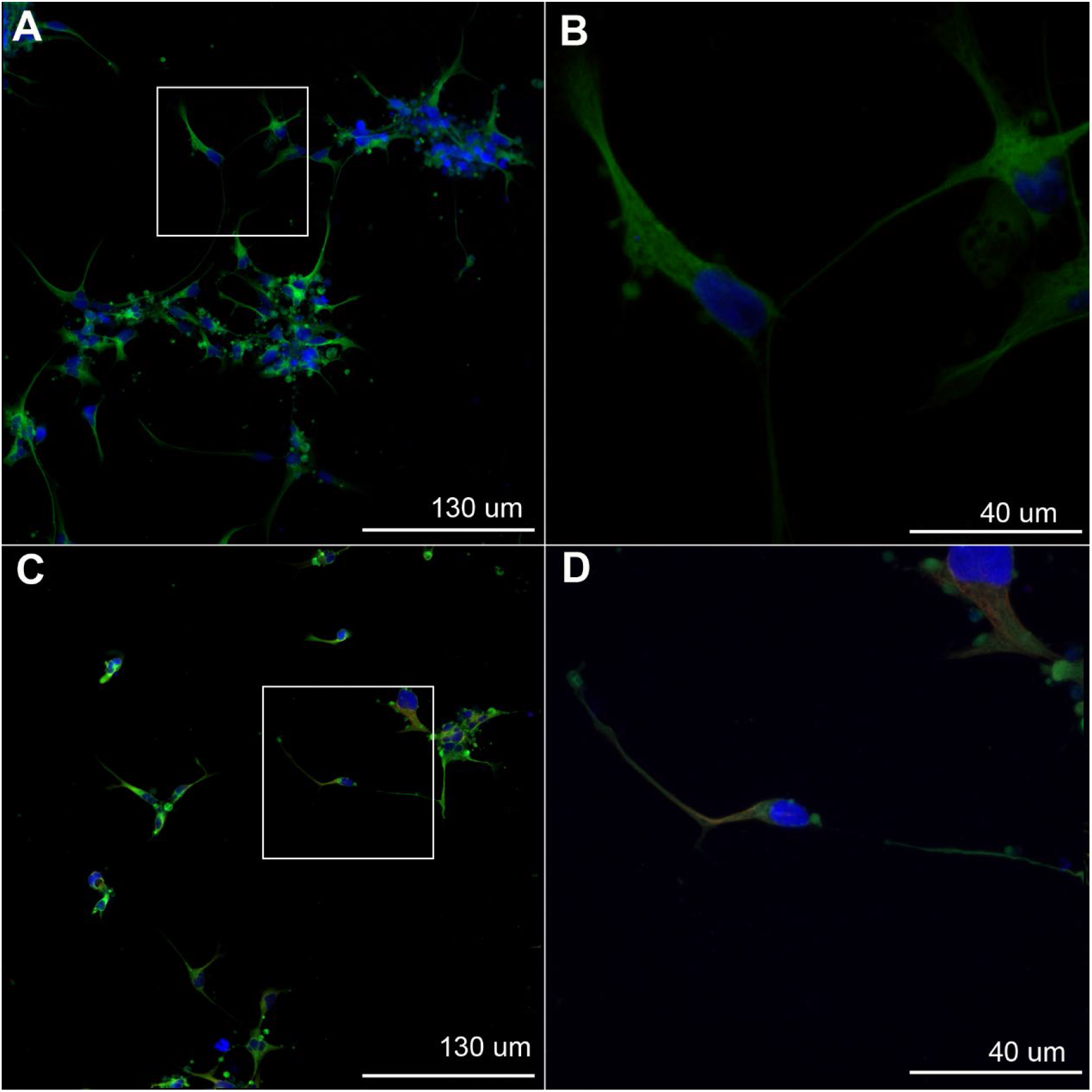
Polyglutamylated Tubulin and βIII-tubulin in Differentiated Neurons (7 day) Fluorescent signals from nuclei (blue), βIII-tubulin (green), and polyglutamylated (red, A, B) or polyglutamylated (red, C,D) tubulin were captured from hNSCs after 7 days of neuronal differentiation in single optical sections at 20x magnification (A, C) and z-series confocal stacks at 63x magnification (projections; B, D). Regions of image capture for higher magnification images are depicted in white boxes in A and C. Image contrast was enhanced to display fluorescent βIII-tubulin signal (75% contrast), and polyglutamylated tubulin signal (93% contrast). Image capture settings and enhancements paralleled those for images of polyglutamylated and acetylated tubulin in basal hNSCs (Fig. 3), and acetylated tubulin in hNSCs after 7 days of neuronal differentiation (Fig. 4).

**Figure 6.**
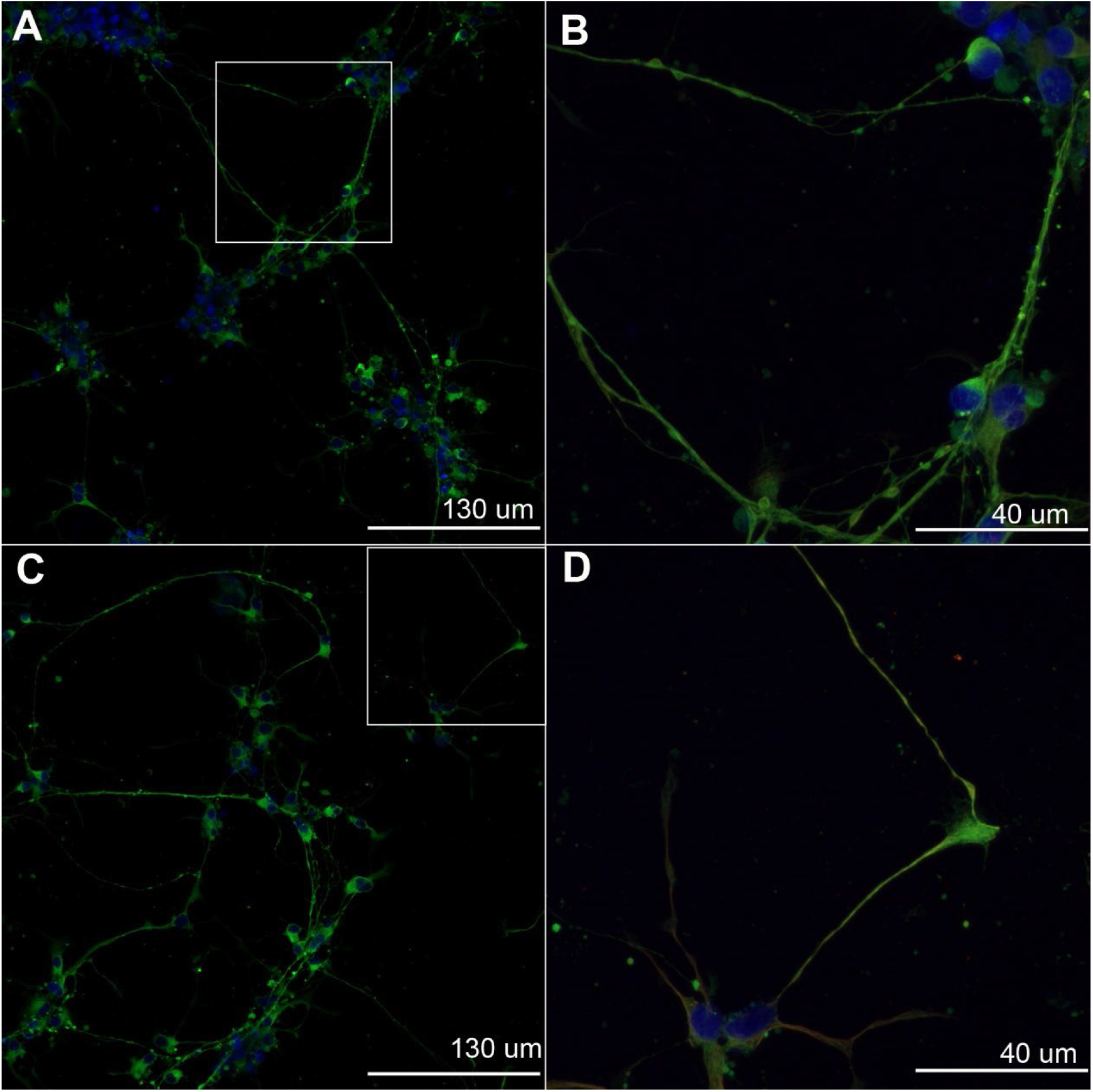
Acetylated Tubulin and βIII-tubulin in Differentiated Neurons (14 day) Fluorescent signals from nuclei (blue), βIII-tubulin (green), and acetylated tubulin (red) were captured from hNSCs after 14 days of neuronal differentiation in single optical sections at 20x magnification (A, C) and z-series confocal stacks at 63x magnification (projections; B, D). Regions of image capture for higher magnification images are depicted in white boxes in A and C. Image contrast was enhanced to display fluorescent βIII-tubulin signal (75% contrast), and acetylated tubulin signal (93% contrast). Image capture settings and enhancements paralleled those for images of polyglutamylated tubulin in hNSCs after 14 days of neuronal differentiation (Fig. 7).

**Figure 7.**
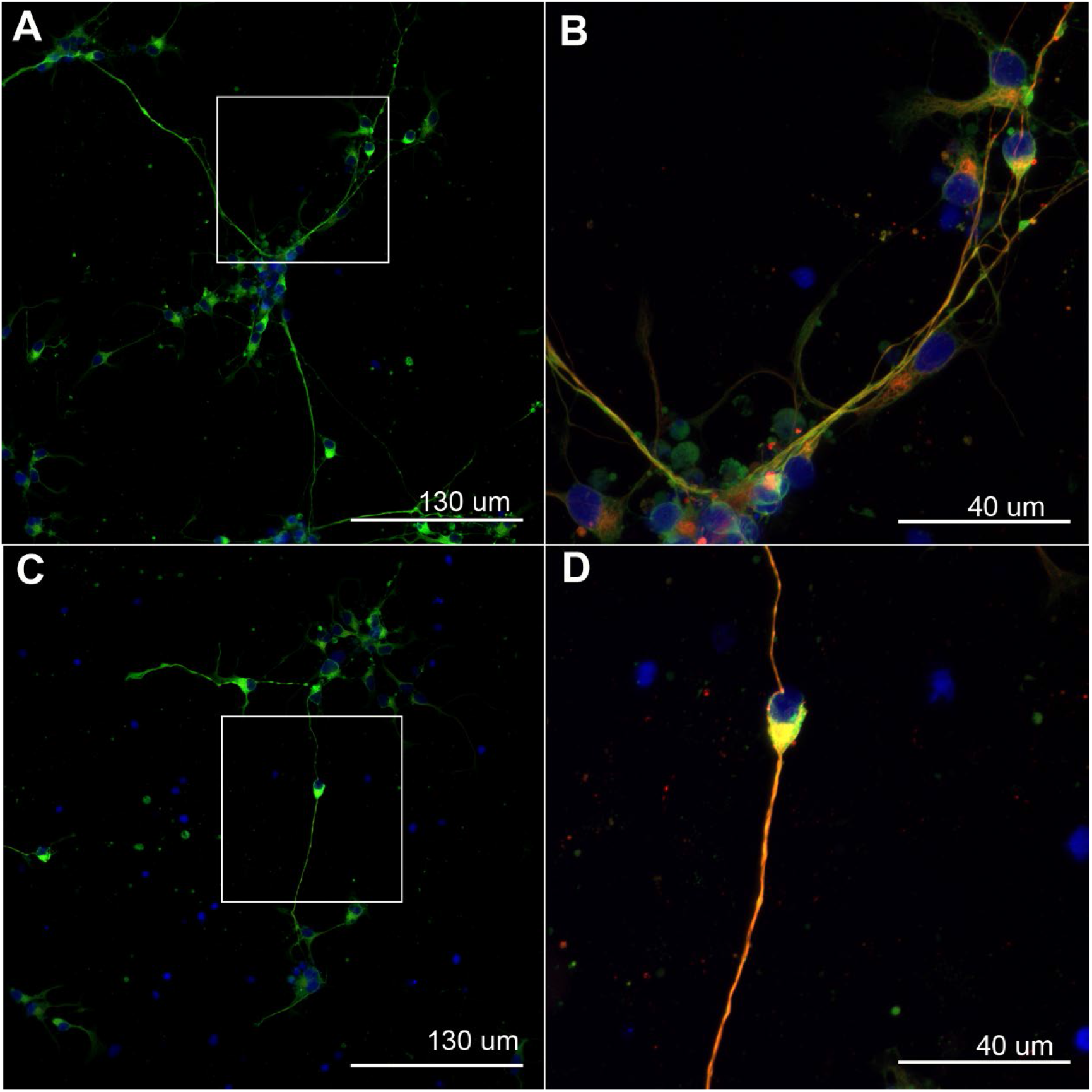
Polyglutamylated Tubulin and βIII-tubulin in Differentiated Neurons (14 day) Fluorescent signals from nuclei (blue), βIII-tubulin (green), and polyglutamylated tubulin (red) were captured from hNSCs after 14 days of neuronal differentiation in single optical sections at 20x magnification (A, C) and z-series confocal stacks at 63x magnification (projections; B, D). Regions of image capture from higher magnification images are depicted in white boxes in A and C. Image contrast was enhanced to display fluorescent βIII-tubulin signal (75% contrast), and polyglutamylated tubulin signal (93% contrast). Image capture settings and enhancements paralleled those for images of acetylated tubulin in hNSCs after 14 days of neuronal differentiation (Fig. 6).

**Figure 8.**
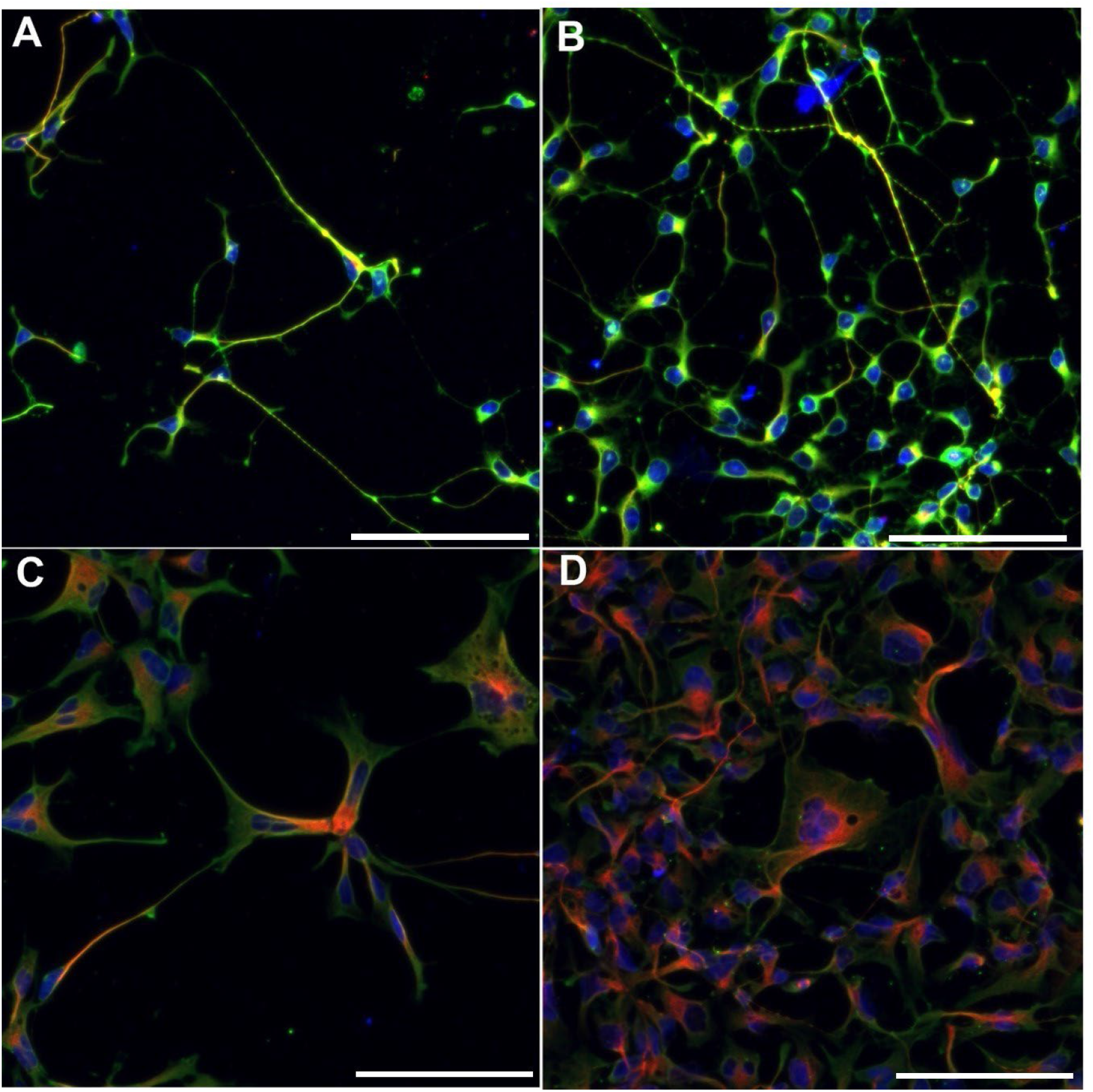
Acetylated Tubulin and βIII-tubulin in Differentiated Neurons (10 day) and Astrocytes. Fluorescent signals from nuclei (blue), βIII-tubulin (green), and acetylated tubulin (red) were captured in z-series confocal stacks (maximum intensity projections displayed) from hNSCs after 10 days of neuronal differentiation (A, B) or 21 days of astroglial differentiation (C,D). Image contrast was not enhanced in the images. Image capture settings paralleled those for images of polyglutamylated tubulin in hNSCs after 10 days of neuronal and 21 days of astroglial differentiation (Fig. 9). Scale bar = 100 μm.

**Figure 9.**
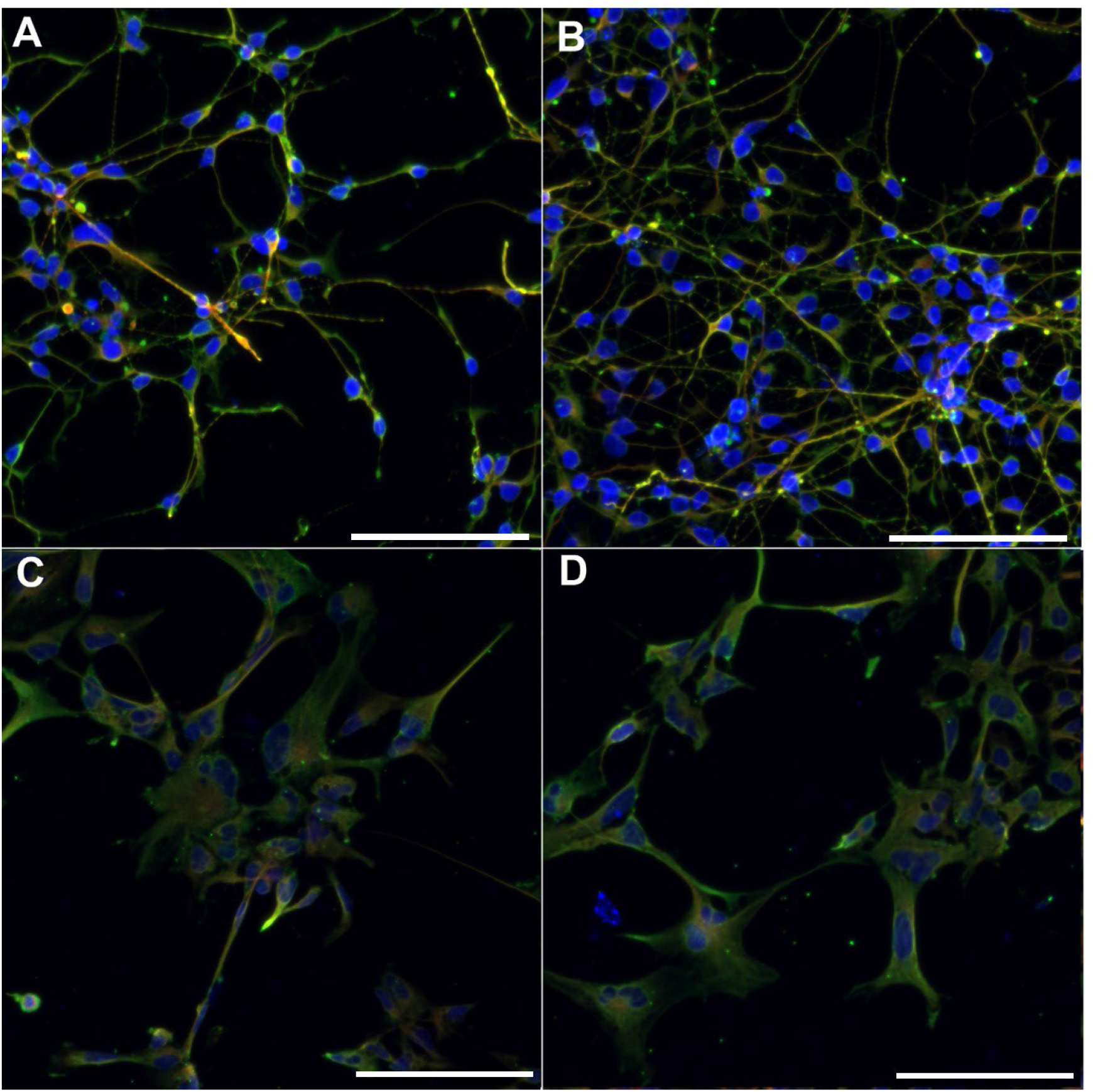
Polyglutamylated Tubulin and βIII-tubulin in Differentiated Neurons (10 day) and Astrocytes. Fluorescent signals from nuclei (blue), βIII-tubulin (green), and polyglutamylated tubulin (red) were captured in z-series confocal stacks (maximum intensity projections displayed) from hNSCs after 10 days of neuronal differentiation (A, B) or 21 days of astroglial differentiation (C,D). Image brightness and contrast were not adjusted. Image capture settings paralleled those for images of acetylated tubulin in hNSCs after 10 days of neuronal and 21 days of astroglial differentiation (Fig. 8). Scale bar = 100 μm.

In basal hNSC cultures, a qualitative assessment revealed that acetylation and polyglutamylation (Fig. 3) were both present, yet at low levels in comparison with differentiated cultures (Figs. 4-9). Polyglutamylation appeared to be more prevalent than acetylation in basal hNSCs (Fig. 3). After seven days of neuronal differentiation, qualitative analysis indicated that acetylation (Fig. 4) was more prevalent than polyglutamylation (Fig. 5). In contrast, after 14 days of neuronal differentiation, polyglutamylation (Fig. 7) appeared more prevalent than acetylation (Fig. 6). After 21 days of astrocyte differentiation, acetylation (Fig. 8) appeared more prevalent than polyglutamylation (Fig. 9).

### 3.3 Quantitation of fluorescence intensity suggests differences in tubulin PTM between media-differentiated neuronal and astroglial hNSC progeny

Mean values for the ratios of PTM to βIII-tubulin were calculated for hNSC progeny after 10 days of neuronal differentiation and 21 days of astroglial differentiation. Acetylation and polyglutamylation were compared within each cell type (neurons, astrocytes). Potential differences in the level of each PTM (acetylation, polyglutamylation) between neurons and astrocytes were also evaluated. The degrees of post-translational modifications of tubulin (acetylation or polyglutamylation) were estimated as a ratio of the sum of fluorescent signal from acetylated or polyglutamylated tubulin to the sum of fluorescent signal from βIII-tubulin in each image.

The mean ratios of acetylated tubulin / βIII-tubulin and polyglutamylated tubulin/ βIII-tubulin were comparable in neurons (∼0.7; Table 2). In astrocytes, a greater divergence was found between the mean ratios of acetylated tubulin/ βIII-tubulin (0.97) and polyglutamylated tubulin/ βIII-tubulin (0.63; p < α = 0.01; Table 2). The mean ratio of polyglutamylated tubulin/ βIII-tubulin was 0.72 in neurons and 0.63 in astrocytes (Table 2). While the mean ratios of acetylated tubulin / βIII-tubulin in neurons (0.73) and astrocytes (0.97) were more divergent (p > α = 0.01; Table 2).

**Table 2.** Statistical comparison of acetylated tubulin/ βIII-tubulin and polyglutamylated tubulin/ βIII-tubulin pixel intensity values in hNSCs after 10 days of neuronal differentiation and 21 days of astrocytic differentiation. Images of neuronal (n=12) and astroglial (n = 5) progeny from sessions 4 and 5, where the magnification and master gain were identical (Table 2), were combined and evaluated for potential differences in posttranslational modifications. Students t-test revealed differences between the two posttranslational tubulin modifications in astrocytes for acetylation and polyglutamylation with p < 0.01, denoted with *.

| **Test 1** | **Cell Type** | **Mean** | **$\sigma$** | **T-value** | **DF** | **p Value** | **Power**<br>**(1 – $\beta$)** |
| --- | --- | --- | --- | --- | --- | --- | --- |
| **Acetylated Tubulin/**<br>**βIII-tubulin** | Neurons | 0.73 | 0.25 | 0.11 | 22 | 0.91 | 0.01 |
| **Polyglutamylated**<br>**Tubulin/ βIII-tubulin** | Neurons | 0.72 | 0.12 |  |  |  |  |
| **Test 2** | **Cell Type** | **Mean** | **$\sigma$** | **T-value** | **D.F.** | **p Value** | **Power**<br>**(1 – $\beta$)** |
| **Acetylated Tubulin/**<br>**βIII-tubulin** | Astrocytes | 0.97 | 0.19 | 3.71 | 8 | 0.006* | 1.00 |
| **Polyglutamylated**<br>**Tubulin/ βIII-tubulin** | Astrocytes | 0.63 | 0.09 |  |  |  |  |
| **Test 3** | **Cell Type** | **Mean** | **$\sigma$** | **T-value** | **D.F.** | **p Value** | **Power**<br>**(1 – $\beta$)** |
| **Acetylated Tubulin/**<br>**βIII-tubulin** | Neurons | 0.73 | 0.25 | 1.95 | 16 | 0.07 | 0.37 |
| **Acetylated Tubulin/**<br>**βIII-tubulin** | Astrocytes | 0.97 | 0.19 |  |  |  |  |
| **Test 4** | **Cell Type** | **Mean** | **$\sigma$** | **T-value** | **D.F.** | **p Value** | **Power**<br>**(1 – $\beta$)** |
| **Polyglutamylated**<br>**Tubulin/ βIII-tubulin** | Neurons | 0.72 | 0.12 | 1.50 | 16 | 0.15 | 0.21 |
| **Polyglutamylated**<br>**Tubulin/ βIII-tubulin** | Astrocytes | 0.63 | 0.09 |  |  |  |  |

## 4. DISCUSSION

Despite the current emphasis on the significance of posttranslational tubulin modifications in the central nervous system, tubulin acetylation and polyglutamylation patterns during the differentiation of neural stem cells have not been thoroughly explored (except see Cao et al., 2017; Sheikh et al. 2021). The expression of βIII-tubulin in neurons is well documented (Moody, Quigg & Frankfurter, 1989; Lee, Rebhun & Frankfurter, 1990). There are relatively fewer demonstrations of βIII-tubulin expression in astrocytes and neural stem cells. However, the presence of βIII-tubulin is becoming established in human neural stem cells as well as plastic cell types and human fetal astrocytes (Katsetos et al., 2001, 2002, 2015; Dráberová et al., 2008; Oikari et al., 2016; Knight & Serrano, 2017a). Astrocytes are diverse and abundant in the CNS (Nedergaard, Ransom & Goldman, 2003; Oberheim et al., 2009; Matyash & Kettenmann, 2010; Stipursky et al., 2012). While the tubulin code has been credited with conferring the diverse array of functional specifications of tubulin, the exploration of potential connections between the tubulin code and astrocyte specialization is in its infancy. This research was an attempt to gain insight into tubulin specifications in human neural stem cells cultured in basal conditions and following differentiation into neurons and astrocytes. Immunocytochemical methods were used to detect βIII-tubulin, acetylated tubulin, and polyglutamylated tubulin in hNSCs.

In accordance with previous reports, we observed the expression of βIII-tubulin in hNSCs in all the culture conditions that were used in this study (Oikari et al., 2016). Quantification and statistical analysis of fluorescence intensity demonstrated that the fluorescence probe intensity ratios for acetylated tubulin / βIII-tubulin and polyglutamylated tubulin/ βIII-tubulin were similar after 10 days of neuronal differentiation. In contrast, after astrocytic differentiation, the ratio of acetylated tubulin/ βIII-tubulin was greater than the ratio of polyglutamylated tubulin/ βIII-tubulin (p < α = 0.01; Table 2). This finding is concordant with previous reports of more acetylation than polyglutamylation in human astrocytes (Knight & Serrano, 2017a).

Given that neurons and basal hNSCs are plated at the same initial seeding density, the neuronal differentiation conditions seem to impact the density of observed cells as seen in phase contast images of basal hNSCs and neuronal progeny (Figs. 1 & 2). This could be due to the lack of cell division that is characteristic of cells that have committed to a neuronal lineage and are in a post-mitotic state (Herrup, 2004; Deneris & Hobert, 2014). Qualitative analysis indicated that acetylated tubulin and polyglutamylated tubulin were also expressed in varying degrees throughout the course of neuronal differentiation. After seven days of differentiation, acetylation appeared more dominant than polyglutamylation in neuronal progeny (Figs. 4 & 5). After 10 days of differentiation, there is no distinct difference between the degrees of acetylation and polyglutamylation in neuronal progeny (Figs. 8 & 9). Finally, after 14 days of neuronal differentiation, polyglutamylation is more prevalent than acetylation in hNSC-derived neurons. This finding is contextualized by the increased levels of tubulin polyglutamylation that are seen in neurodegeneration, which increases in prevalence in proportion to age (Zempel et al., 2013; Vu et al., 2017). Moreover, a putative role for polyglutamylated tubulin in neurodegenerative disorders is through the regulation of endoplasmic reticulum morphology (Zheng et al., 2022).

Whereas the link between polyglutamylation and neurodegeneration is being established, there is still relatively little information about the role of tubulin acetylation and polyglutamylation in the specification of neural cell fate (Moutin et al., 2021). The changing profile of tubulin modifications during neuronal development has been established in mouse neurons (Audebert et al., 1994; Ikegami et al., 2006). However, we have yet to understand the precise role of PTMs in the process of differentiation, particularly of human neural cell types. Acetylation and polyglutamylation have a demonstrated role in cell fate specification overall, as seen in the pre-implantation embryo and during mouse development (Houliston & Maro, 1989; Zhang et al., 2008). Recent studies have also established the role of acetylation in neuron development and morphogenesis, particularly with respect to axon growth and branching (Dan Wei et al., 2018; Saunders et al., 2022). Our findings add to this body of literature and advance the understanding of tubulin PTMs in human neurons and astrocytes during development.

In summary, these results are a preview of the tubulin modification patterns that facilitate the specification of distinct identities among neural cell types. Our findings offer insight into the complex specialization patterns conferred by tubulin PTMs in the specification of neural cell fate and the overarching process of morphogenesis. Outcomes from this study emphasize the need to explore how tubulin specifications contribute to neural cell identity during stem cell differentiation as well as the role of PMTs in glial functional diversity (Schiweck, Eickholt & Murk, 2018). Moreover, these findings present hNSCs and their progeny as a potential model for the complex processes of morphogenesis and neurodegeneration.

## Abbreviations

β: beta
BSA: bovine serum albumin
hESC: human embryonic stem cell
hNSC: human neural stem cell
IgG: immunoglobulin
MAP: microtubule associated protein
PBS: phosphate buffered saline
PBST: phosphate buffered saline with 0.1% TWEEN® 20
RRID: Research Resource Identifiers
PTM: Post-translational modifications

## Author Contributions

**V. Bleu Knight** conceived and designed the experiments, performed the experiments, analyzed the data, interpreted results, prepared the first draft of the manuscript, prepared all figures and/or tables, reviewed drafts of the paper.

**Manasi P. Jogalekar** performed experiments and reviewed drafts of the paper.

**Elba E. Serrano** conceived and designed experiments, analyzed the data, interpreted results, contributed reagents/materials/analysis tools, co-wrote the paper, reviewed drafts of the paper, supervised project, provided oversight for compliance with institutional agency procedures for research integrity.

## Electronic supplementary material (NONE)

## Acknowledgement

We would like to thank Dr. Armando Varela-Ramirez of the BBRC-CSIC Facility for assistance with confocal imaging.

## REFERENCES

Audebert S, Desbruyères E, Gruszczynski C, Koulakoff a, Gros F, Denoulet P, Eddé B. 1993. Reversible polyglutamylation of alpha- and beta-tubulin and microtubule dynamics in mouse brain neurons. Molecular biology of the cell 4:615–626. DOI: 10.1091/mbc.4.6.615.

Audebert S, Koulakoff a, Berwald-Netter Y, Gros F, Denoulet P, Eddé B. 1994. Developmental regulation of polyglutamylated alpha- and beta-tubulin in mouse brain neurons. Journal of cell science 107 (Pt 8:2313–22. DOI: 10.1016/j.ctrv.2012.07.005.

Bandrowski A, Brush M, Grethe JS, Haendel MA, Kennedy DN, Hill S, Hof PR, Martone ME, Pols M, Tan SC, Washington N, Zudilova-Seinstra E, Vasilevsky N. 2016. The Resource Identification Initiative: A cultural shift in publishing. Brain and Behavior 6:1–14. DOI: 10.1002/brb3.417.

Bär J, Popp Y, Bucher M, Mikhaylova M. 2022. Direct and indirect effects of tubulin post-translational modifications on microtubule stability: Insights and regulations. Biochimica et Biophysica Acta (BBA) - Molecular Cell Research 1869:119241. DOI: 10.1016/j.bbamcr.2022.119241.

Bodakuntla S, Yuan X, Genova M, Gadadhar S, Leboucher S, Birling M, Klein D, Martini R, Janke C, Magiera MM. 2021. Distinct roles of α- and β-tubulin polyglutamylation in controlling axonal transport and in neurodegeneration. The EMBO Journal 40. DOI: 10.15252/embj.2021108498.

Cambray-Deakin MA, Burgoyne RD. 1987. Posttranslational modifications of α-tubulin: Acetylated and detyrosinated forms in axons of rat cerebellum. Journal of Cell Biology 104:1569–1574. DOI: 10.1083/jcb.104.6.1569.

Casale CH, Previtali G, Barra HS. 2003. Involvement of acetylated tubulin in the regulation of Na+,K+-ATPase activity in cultured astrocytes. FEBS Letters 534:115–118. DOI: 10.1016/S0014-5793(02)03802-4.

Chakraborti S, Natarajan K, Curiel J, Janke C, Liu J. 2016. The emerging role of the tubulin code: from the tubulin molecule to neuronal function and disease. Cytoskeleton (Hoboken, N.J.). DOI: 10.1002/cm.21290.

Dan Wei, Gao N, Li L, Zhu J-X, Diao L, Huang J, Han Q-J, Wang S, Xue H, Wang Q, Wu Q-F, Zhang X, Bao L. 2018. α-Tubulin Acetylation Restricts Axon Overbranching by Dampening Microtubule Plus-End Dynamics in Neurons. Cerebral Cortex 28:3332–3346. DOI: 10.1093/cercor/bhx225.

Delépine C, Meziane H, Nectoux J, Opitz M, Smith AB, Ballatore C, Saillour Y, Bennaceur-Griscelli A, Chang Q, Williams EC, Dahan M, Duboin A, Billuart P, Herault Y, Bienvenu T. 2016. Altered microtubule dynamics and vesicular transport in mouse and human MeCP2-deficient astrocytes. Human Molecular Genetics 25:146–157. DOI: 10.1093/hmg/ddv464.

Deneris ES, Hobert O. 2014. Maintenance of postmitotic neuronal cell identity. Nature Neuroscience 17:899–907. DOI: 10.1038/nn.3731.

Dráberová E, Del Valle L, Gordon J, Marková V, Smejkalová B, Bertrand L, de Chadarévian J-P, Agamanolis DP, Legido A, Khalili K, Dráber P, Katsetos CD. 2008. Class III beta-tubulin is constitutively coexpressed with glial fibrillary acidic protein and nestin in midgestational human fetal astrocytes: implications for phenotypic identity. Journal of neuropathology and experimental neurology 67:341–54. DOI: 10.1097/NEN.0b013e31816a686d.

Faul F, Erdfelder E, Buchner A, Lang A-G. 2009. Statistical power analyses using G*Power 3.1: Tests for correlation and regression analyses. Behavior Research Methods 41:1149–1160. DOI: 10.3758/BRM.41.4.1149.

Gadau SD. 2015. Detection, Distribution and Amount of Posttranslational α -Tubulin Modifications in Immortalized Rat Schwann Cells. :255–265. DOI: 10.12871/00039829201542.

Herrup K. 2004. Divide and Die: Cell Cycle Events as Triggers of Nerve Cell Death. Journal of Neuroscience 24:9232–9239. DOI: 10.1523/JNEUROSCI.3347-04.2004.

Houliston E, Maro B. 1989. Posttranslational modification of distinct microtubule subpopulations during cell polarization and differentiation in the mouse preimplantation embryo. Journal of Cell Biology 108:543–551. DOI: 10.1083/jcb.108.2.543.

Ikegami K, Mukai M, Tsuchida JI, Heier RL, MacGregor GR, Setou M. 2006. TTLL7 is a mammalian ??-tubulin polyglutamylase required for growth of MAP2-positive neurites. Journal of Biological Chemistry 281:30707–30716. DOI: 10.1074/jbc.M603984200.

Janke C, Kneussel M. 2010. Tubulin post-translational modifications: Encoding functions on the neuronal microtubule cytoskeleton. Trends in Neurosciences 33:362–372. DOI: 10.1016/j.tins.2010.05.001.

Janke C, Magiera MM. 2020. The tubulin code and its role in controlling microtubule properties and functions. Nature Reviews Molecular Cell Biology 21:307–326. DOI: 10.1038/s41580-020-0214-3.

Katsetos CD, Reginato MJ, Baas PW, D’Agostino L, Legido A, Tuszyn Ski JA, Dráberová E, Dráber P. 2015. Emerging microtubule targets in glioma therapy. Seminars in Pediatric Neurology 22:49–72. DOI: 10.1016/j.spen.2015.03.009.

Katsetos CD, Del Valle L, Geddes JF, Aldape K, Boyd JC, Legido A, Khalili K, Perentes E, Mörk SJ. 2002. Localization of the neuronal class III beta-tubulin in oligodendrogliomas: comparison with Ki-67 proliferative index and 1p/19q status. Journal of Neuropathology and Experimental Neurology 61:307–320.

Katsetos CD, Del Valle L, Geddes JF, Assimakopoulou M, Legido A, Boyd JC, Balin B, Parikh N a, Maraziotis T, de Chadarevian JP, Varakis JN, Matsas R, Spano A, Frankfurter A, Herman MM, Khalili K. 2001. Aberrant localization of the neuronal class III beta-tubulin in astrocytomas. Archives of pathology & laboratory medicine 125:613–24. DOI: 10.1043/0003-9985(2001)125<0613:ALOTNC>2.0.CO;2.

Knight VB, Serrano EE. 2017a. Post-translational Tubulin Modifications in Human Astrocyte Cultures. Neurochemical Research. DOI: 10.1007/s11064-017-2290-0.

Knight VB, Serrano EE. 2017b. Hydrogel scaffolds promote neural gene expression and structural reorganization in human astrocyte cultures. PeerJ. DOI: 10.7717/peerj.2829.

Knight VB, Serrano EE. 2017c. RNA sequencing analysis of neural cell lines: impact of normalization and technical replication. In: Rojas I, Ortuño F eds. Bioinformatics and Biomedical Engineering. Granada: Springer, 457–468.

Landis SC, Amara SG, Asadullah K, Austin CP, Blumenstein R, Bradley EW, Crystal RG, Darnell RB, Ferrante RJ, Fillit H, Finkelstein R, Fisher M, Gendelman HE, Golub RM, Goudreau JL, Gross RA, Gubitz AK, Hesterlee SE, Howells DW, Huguenard J, Kelner K, Koroshetz W, Krainc D, Lazic SE, Levine MS, Macleod MR, McCall JM, Moxley RT, Narasimhan K, Noble LJ, Perrin S, Porter JD, Steward O, Unger E, Utz U, Silberberg SD. 2012. A call for transparent reporting to optimize the predictive value of preclinical research. Nature 490:187–191. DOI: 10.1038/nature11556.

Lee MK, Rebhun LI, Frankfurter a. 1990. Posttranslational modification of class III beta-tubulin. Proceedings of the National Academy of Sciences of the United States of America 87:7195–7199. DOI: 10.1073/pnas.87.18.7195.

Lessard D V., Zinder OJ, Hotta T, Verhey KJ, Ohi R, Berger CL. 2019. Polyglutamylation of tubulin’s C-terminal tail controls pausing and motility of kinesin-3 family member KIF1A. Journal of Biological Chemistry 294:6353–6363. DOI: 10.1074/jbc.RA118.005765.

Li L, Yang XJ. 2015. Tubulin acetylation: Responsible enzymes, biological functions and human diseases. Cellular and Molecular Life Sciences 72:4237–4255. DOI: 10.1007/s00018-015-2000-5.

Matyash V, Kettenmann H. 2010. Heterogeneity in astrocyte morphology and physiology. Brain Research Reviews 63:2–10. DOI: 10.1016/j.brainresrev.2009.12.001.

Moody SA, Quigg MS, Frankfurter A. 1989. Development of the peripheral trigeminal system in the chick revealed by an isotype-specific anti-beta-tubulin monoclonal antibody. The Journal of Comparative Neurology 279:567–580. DOI: 10.1002/cne.902790406.

Moutin M, Bosc C, Peris L, Andrieux A. 2021. Tubulin post-translational modifications control neuronal development and functions. Developmental Neurobiology 81:253–272. DOI: 10.1002/dneu.22774.

Nedergaard M, Ransom B, Goldman SA. 2003. New roles for astrocytes: Redefining the functional architecture of the brain. Trends in Neurosciences 26:523–530. DOI: 10.1016/j.tins.2003.08.008.

Oberheim NA, Takano T, Han X, He W, Lin JH, Wang F, Xu Q, Wyatt JD, Pilcher W, Ojemann JG, Ransom BR, Goldman SA, Nedergaard M. 2009. Uniquely hominid features of adult human astrocytes. Journal of Neuroscience 29:3276–3287. DOI: 10.1523/jneurosci.4707-08.2009.

Oikari LE, Okolicsanyi RK, Qin A, Yu C, Grif LR, Haupt LM. 2016. Cell surface heparan sulfate proteoglycans as novel markers of human neural stem cell fate determination. 16:92–104. DOI: 10.1016/j.scr.2015.12.011.

Park JH, Roll-Mecak A. 2018. The tubulin code in neuronal polarity. Current Opinion in Neurobiology 51:95–102. DOI: 10.1016/j.conb.2018.03.001.

Pau G, Fuchs F, Sklyar O, Boutros M, Huber W. 2010. EBImage-an R package for image processing with applications to cellular phenotypes. Bioinformatics 26:979–981. DOI: 10.1093/bioinformatics/btq046.

Pellegrini L, Wetzel A, Grannó S, Heaton G, Harvey K. 2017. Back to the tubule: microtubule dynamics in Parkinson’s disease. Cellular and Molecular Life Sciences 74:409–434. DOI: 10.1007/s00018-016-2351-6.

Roll-Mecak A. 2020. The Tubulin Code in Microtubule Dynamics and Information Encoding. Developmental Cell 54:7–20. DOI: 10.1016/j.devcel.2020.06.008.

Romaniello R, Arrigoni F, Bassi MT, Borgatti R. 2015. Mutations in ??- and ??-tubulin encoding genes: Implications in brain malformations. Brain and Development 37:273–280. DOI: 10.1016/j.braindev.2014.06.002.

Ruse CI, Chin HG, Pradhan S. 2022. Polyglutamylation: biology and analysis. Amino Acids 54:529–542. DOI: 10.1007/s00726-022-03146-4.

Santiago-Mujika E, Luthi-Carter R, Giorgini F, Kalaria RN, Mukaetova-Ladinska EB. 2021. Tubulin and Tubulin Posttranslational Modifications in Alzheimer’s Disease and Vascular Dementia. Frontiers in Aging Neuroscience 13. DOI: 10.3389/fnagi.2021.730107.

Saragoni L, Hernández P, Maccioni RB. 2000. Differential association of tau with subsets of microtubules containing posttranslationally-modified tubulin variants in neuroblastoma cells. Neurochemical Research 25:59–70. DOI: 10.1023/A:1007587315630.

Saunders HAJ, Johnson-Schlitz DM, Jenkins B V., Volkert PJ, Yang SZ, Wildonger J. 2022. Acetylated α-tubulin K394 regulates microtubule stability to shape the growth of axon terminals. Current Biology 32:614-630.e5. DOI: 10.1016/j.cub.2021.12.012.

Schiweck J, Eickholt BJ, Murk K. 2018. Important Shapeshifter: Mechanisms Allowing Astrocytes to Respond to the Changing Nervous System During Development, Injury and Disease. Frontiers in Cellular Neuroscience 12. DOI: 10.3389/fncel.2018.00261.

Seetharaman S, Vianay B, Roca V, Farrugia AJ, De Pascalis C, Boëda B, Dingli F, Loew D, Vassilopoulos S, Bershadsky A, Théry M, Etienne-Manneville S. 2022. Microtubules tune mechanosensitive cell responses. Nature Materials 21:366–377. DOI: 10.1038/s41563-021-01108-x.

Song Y, Brady ST. 2015. Post-translational modifications of tubulin: Pathways to functional diversity of microtubules. Trends in Cell Biology 25:125–136. DOI: 10.1016/j.tcb.2014.10.004.

Stipursky J, De Sampaio e Spohr TCL, Sousa VO, Gomes FCA. 2012. Neuron-astroglial interactions in cell-fate commitment and maturation in the central nervous system. Neurochemical Research 37:2402–2418. DOI: 10.1007/s11064-012-0798-x.

Tischfield MA, Cederquist GY, Gupta ML, Engle EC. 2011. Phenotypic spectrum of the tubulin-related disorders and functional implications of disease-causing mutations. Current Opinion in Genetics and Development 21:286–294. DOI: 10.1016/j.gde.2011.01.003.

Vu HT, Akatsu H, Hashizume Y, Setou M, Ikegami K. 2017. Increase in α -tubulin modifications in the neuronal processes of hippocampal neurons in both kainic acid-induced epileptic seizure and Alzheimer ‘ s disease. Nature Publishing Group:1–14. DOI: 10.1038/srep40205.

Wu H-Y, Rong Y, Bansal PK, Wei P, Guo H, Morgan JI. 2022. TTLL1 and TTLL4 polyglutamylases are required for the neurodegenerative phenotypes in pcd mice. PLOS Genetics 18:e1010144. DOI: 10.1371/journal.pgen.1010144.

Yoshiyama Y, Zhang B, Bruce J, Trojanowski JQ, Lee VM-Y. 2003. Reduction of detyrosinated microtubules and Golgi fragmentation are linked to tau-induced degeneration in astrocytes. The Journal of neuroscience : the official journal of the Society for Neuroscience 23:10662–10671. DOI: 23/33/10662 [pii].

Youn GS, Ju SM, Choi SY, Park J. 2015. HDAC6 mediates HIV-1 tat-induced proinflammatory responses by regulating MAPK-NF-kappaB/AP-1 pathways in astrocytes. Glia 63:1953–1965. DOI: 10.1002/glia.22865.

Zempel H, Luedtke J, Kumar Y, Biernat J, Dawson H, Mandelkow E, Mandelkow E-M. 2013. Amyloid-β oligomers induce synaptic damage via Tau-dependent microtubule severing by TTLL6 and spastin. The EMBO Journal 32:2920–2937. DOI: 10.1038/emboj.2013.207.

Zhang Y, Kwon S, Yamaguchi T, Cubizolles F, Rousseaux S, Kneissel M, Cao C, Li N, Cheng H-L, Chua K, Lombard D, Mizeracki A, Matthias G, Alt FW, Khochbin S, Matthias P. 2008. Mice Lacking Histone Deacetylase 6 Have Hyperacetylated Tubulin but Are Viable and Develop Normally. Molecular and Cellular Biology 28:1688–1701. DOI: 10.1128/MCB.01154-06.

Zhang F, Su B, Wang C, Siedlak SL, Mondragon-Rodriguez S, Lee H, Wang X, Perry G, Zhu X. 2015. Posttranslational modifications of α-tubulin in alzheimer disease. Translational Neurodegeneration 4:9. DOI: 10.1186/s40035-015-0030-4.

Zheng P, Obara CJ, Szczesna E, Nixon-Abell J, Mahalingan KK, Roll-Mecak A, Lippincott-Schwartz J, Blackstone C. 2022. ER proteins decipher the tubulin code to regulate organelle distribution. Nature 601:132–138. DOI: 10.1038/s41586-021-04204-9.

